# Aberrant ventral dentate gyrus structure and function in individuals susceptible to post-traumatic stress disorder

**DOI:** 10.1101/2020.10.01.321893

**Authors:** Bart C.J. Dirven, Dewi van der Geugten, Miranda van Bodegom, Leonie Madder, Laura van Agen, Judith R. Homberg, Tamas Kozicz, Marloes J.A.G. Henckens

## Abstract

Post-traumatic stress disorder (PTSD) is a psychiatric disorder vulnerable individuals can develop following a traumatic event, whereas others are resilient. Enhanced insight into the mechanistic underpinnings contributing to these inter-individual differences in PTSD susceptibility is key to improved treatment and prevention. Aberrant function of the hippocampal dentate gyrus (DG) may contribute to its psychopathology, with the dorsal DG potentially encoding trauma memory generalization and the ventral DG anxiety. Using a mouse model, we investigated the association between deviant DG structure and function and susceptibility to develop PTSD-like symptoms following trauma. Mice were exposed to a traumatic event (unpredictable, inescapable foot shocks) and tested for PTSD symptomatology following recovery. In three independent experiments, DG neuronal morphology, synaptic protein gene expression and neuronal activity during trauma encoding and recall were assessed. Behaviorally, PTSD-like animals displayed some increased anxiety-like behavior already prior to trauma, increased novelty-induced freezing, but no clear differences in remote trauma memory recall. Comparison of the ventral DG of PTSD-like vs resilient mice revealed lower spine density, reduced expression of the postsynaptic protein homer 1b/c gene, a larger population of neurons active during trauma encoding and a greater presence of somatostatin neurons to be associated with PTSD susceptibility. In contrast, the dorsal DG of PTSD-like animals did not differ in terms of spine density or gene expression, but displayed more active neurons during trauma encoding and a lower amount of somatostatin neurons. These data propose a critical role for -mainly the ventral-DG in establishing symptomatology addressed in this PTSD model.

## INTRODUCTION

Posttraumatic stress disorder (PTSD) is a debilitating disorder one can develop after exposure to a traumatic event. PTSD patients experience excessive arousal, hypervigilance, exaggerated startle responses, and insomnia (DSM-V^1^), which severely impact quality of life. Moreover, one of the hallmark features of PTSD is the re-experiencing of the trauma by flashbacks, spontaneous recollections, and recurrent nightmares of the trauma^2^. However, whereas over 80% of individuals ever experience a traumatic event, only a relatively small fraction (∼15%) will develop PTSD^3,4^. Understanding the neural basis of this inter-individual variability in PTSD susceptibility is critical for understanding PTSD psychopathology^5^, and likely holds unique insights for optimized treatment and even prevention.

An over-generalization of fear to safe, non-trauma related situations is thought to contribute to PTSD psychopathology^6,7^, but the exact underlying mechanisms remain unclear. Previous research has implicated aberrant function of the hippocampal dentate gyrus (DG) in fear generalization, impairing hippocampal pattern separation^8^, a process resolving interference in encoding and retrieving similar experiences^9-14^. Robust lateral inhibition of DG granule cells by inhibitory interneurons in the hilar region^11,15^ ensures the sparse activation necessary for efficient pattern separation, with a prominent role for somatostatin-expressing (SOM) interneurons^16^. However, pattern separation capacity has mainly been attributed to the dorsal DG (dDG), whereas the ventral DG (vDG) seems to be more involved in affective processing^17-20^. Activity in the vDG is associated with anxiety^18,21-23^, the return of extinguished fear^24^, and mediating the anxiolytic effects of antidepressant treatment^25-27^. These findings suggest that the dDG might contribute to PTSD symptomatology by impaired pattern separation processes, whereas the vDG might be implicated by mediating increased anxiety. Supporting a role for aberrant overall DG function in PTSD, patients show poor performance on a memory task testing pattern separation^28^, as well as a smaller DG volume^29,30^, which correlates with PTSD symptom severity^29^. Rodent work has added to these findings by showing reduced dendritic complexity and spine density in the DG of animals most sensitive to a trauma^31-34^, and a reduced expression of DG synaptic proteins^35^. Importantly, most of these studies did not investigate the DG function and structure across its dorsal-ventral axis, making that its exact deviations in the PTSD-affected brain are still largely unknown.

Here, we set out to investigate potential DG abnormalities in a PTSD model in mice, in which mice were first exposed to a traumatic event (foot shocks) and then behaviorally tested for PTSD-like symptomatology, dissociating PTSD-like from resilient mice. To understand the neurobiological basis of PTSD susceptibility, we compared dorsal and ventral DG structure (neuronal morphology and synaptic protein gene expression) and function (activity during trauma memory encoding and remote recall) between groups. Moreover, behavioral readouts of anxiety and fear generalization were assessed prior to or immediately following trauma exposure, to obtain insights into important predictors of later PTSD-like symptoms.

## MATERIALS & METHODS

### Animals

The study consisted of three separate experiments: experiment 1 (n=24) to assess DG spine density, experiment 2 (n=48) for assessing DG gene expression, and experiment 3 (n=40) to assess DG neuronal activity. For experiments 1-2, C57BL/6J mice (Charles River, France) were used. For experiment 3, heterozygote male FosCreER^T2^ (B6.129(Cg)-*Fos*^*tm1*.*1(cre/ERT2)Luo*^/J, 021882, Jackson Laboratory) and homozygote female ROSA mice (B6.Cg-*Gt(ROSA)26Sor*^*tm9(CAG-tdTomato)Hze*^/J, 007909, Jackson Laboratory) were crossed to generate heterozygote offspring (referred to as FosTRAP^36^). Based on sex differences in stress sensitivity^37,38^, and the fact that the used PTSD model has only been validated in males^39,40^, only male mice were used. Mice were group housed (3-4 mice per cage) in individually ventilated cages on a reverse 12 hour (9.00-21.00h) dark/light cycle. Food and water were provided *ad libitum*. All behavioral testing was performed at least 4 hours into the animals’ active phase (i.e., the dark). The experimental protocols were in line with international guidelines, the Care and Use of Mammals in neuroscience and Behavioural Research (National Research Council 2003), the principles of laboratory animal care, as well as the Dutch law concerning animal welfare and approved by the Central Committee for Animal Experiments, Den Haag, The Netherlands.

### PTSD Model

All mice were exposed to a PTSD induction model (Fig. S1) as described before^39,40^. On day 1, mice were exposed to a traumatic event, i.e., the exposure to 14x 1 s 1 mA foot shocks at variable intervals for an 85 min session in a certain context A. On day 2, 21 hours post-trauma, mice were subjected to a subsequent trigger, i.e., 5x 1 s 0.7 mA foot shocks at a fixed 1-min interval for 5 min session, in a different context (context B). Mice were videotaped during trigger exposure, and videos analyzed for freezing behavior by a researcher blind to the experimental group, to assess novelty-induced anxiety as well as shock-induced fear. Mice were allowed to recover, and at a week post-trauma tested for their behavioral response by assessing PTSD-related behavior; impaired risk assessment (dark-light transfer test), increased anxiety (marble burying), hypervigilance (acoustic startle), impaired sensorimotor gaiting (pre-pulse inhibition), and disturbed circadian rhythm (locomotor activity during the light phase)^40^. See the Supplementary Materials for further details.

#### Behavioral Categorization

In order to categorize mice as either PTSD-like or resilient, mouse behavior on each of the tests was sorted and the 25% of mice who had the lowest values were attributed 3 points for percentage risk assessment, 3 points for latency to peak startle amplitude, and 2 points for percentage PPI. Similarly, the 25% of mice showing the highest values were attributed 1 point for light locomotor activity and marble burying^39^. Points for each test were determined by factor analysis as described before^40^. The points per animal were tallied to generate an overall PTSD symptom score, and mice that had totals of 5 or more points (necessitating extreme behavior in multiple tests) were termed PTSD-like. Only mice that had zero points were termed resilient.

### Experiment 1: DG Neuronal Morphology

#### Sacrifice

Mice were subjected to the PTSD model and sacrificed on day 23 under anesthesia (5% isoflurane inhalation followed by i.p. injection with 200 μl pentobarbital) by perfusion with phosphate-buffered saline (PBS) followed by phosphate-buffered 4% paraformaldehyde (PFA). The brains were surgically removed and post-fixed for 24 hours in 4% PFA, after which they were transferred to 0.1 M PBS with 0.01% sodium azide and stored at 4 °C.

#### Golgi Staining

Brains of PTSD-like (n=4) and resilient (n=5) mice were processed for rapid Golgi-Cox staining (FD Rapid GolgiStain™ FDNeurotechnologies, Inc. Ellicott City, MD, USA) to examine the neuronal morphology of dorsal and ventral DG granule cells. Briefly, brain tissue was immersed in a solution consisting of a mixture of mercury chloride, potassium dichromate and potassium chromate, and kept in the dark for 14 days. After impregnation, the tissue was transferred into a cryoprotectant solution for 72 hours, and cut into 140 μm coronal sections using a freezing sliding microtome (Microtom HM440E, GMI Inc., Ramsey, MN, USA), and mounted on gelatin-coated slides. The sections were left to air-dry for 3 days before the sections were washed twice for 4 min in dH_2_O, and incubated for 10 min in an ammonium hydroxide solution. Then, the sections were 2x washed in dH_2_O for 4 min, placed for 4 min in ascending concentrations of ethanol (70%, 95% and four times 100%) and washed 3x 4 min in xylene for clearance. Then, sections were cover slipped with PermountTM mounting medium (Fisher Chemicals, Leicestershire, UK) and dried for 1 week until further processing for analysis. Details about the neuronal reconstruction and spine density analysis are given in the Supplementary Materials.

### Experiment 2: DG Gene Expression

#### Design and sacrifice

Mice in this experiment were first tested in the Open Field and Elevated Plus Maze tests for assessing pre-trauma anxiety (Supplementary Materials). Additionally, the mice were exposed to two functional neuroimaging sessions (7 days prior to trauma induction and 20 days post-trauma) in an 11.7 T BioSpec Avance III small animal MR system (Bruker BioSpin), while anesthetized by 0.5% inhalation isoflurane and subcutaneous infusion of medetomidine (Dexdomitor, Pfizer, 0.1 mg/kg/h^41^). These data are, however, beyond the scope of the present study. Mice were sacrificed on day 28 by rapid decapitation, and brains were surgically removed, quickly frozen on dry ice, and stored at −80 °C until further processing.

#### Isolation of target tissue

Brains of PTSD-like (n=9) and resilient (n=12) mice from experiment 2 were sliced into 300 μm coronal slices on a Leica CM3050 S Research Cryostat (Leica Biosystems, Amsterdam, the Netherlands), with a chamber temperature of −12 °C and an object temperature of −10 °C, after which regions of interest were punched out. dDG punches were taken bilaterally with a 0.5 mm diameter hollow needle from three subsequent slices (Bregma −1.70:−2.30 mm), for a total of six punches per subregion. Similarly, six 0.75 mm diameter punches were taken from the vDG (Bregma −2.80:−3.40 mm).

#### RNA extraction and cDNA synthesis

RNA was extracted from the isolated tissue using the AllPrep DNA/RNA Mini Kit (QIAGEN, Venlo, the Netherlands), after which cDNA was synthesized using the SensiFAST(tm) cDNA Synthesis Kit (Bioline, Taunton, MA, USA).

#### Quantitative PCR

Gene expression was compared in dorsal and ventral DG of PTSD-like and resilient mice using qPCR. We chose to look specifically at pre- and postsynaptic markers to tie into the spine density measurements, and added a spine-localized immediate early gene and a neuronal marker to be able to relate gene expression to the amount of neuronal material included in the punch. Assays included genes encoding synapsin I (*Syn1*) and synaptophysin (*Syp*), both present in synaptic vesicles^42^; postsynaptic density-95 (*Psd-95*), encoding a postsynaptic membrane protein^43^; homer1b/c (*Homer1* splice variant), an postsynaptic density scaffolding protein involved in glutamate receptor transporter protein availability; homer1a (*Homer1* splice variant), an immediate early gene and shown to be in direct competition with the longer transcript homer1b/c^44^; and neurofilament H (*Nefh*), a neuronal marker^35^. Hypoxanthine-guanine phosphoribosyl transferase (*Hprt*) and cytochrome c1 (*Cyc1*) were chosen as housekeeping genes. Details are given in the Supplementary Materials.

### Experiment 3: DG Neuronal Activity

#### Design and sacrifice

Mice in this experiment were injected with tamoxifen solution 7 hours prior to trauma exposure to label trauma-related neuronal activity by the induction of tdTomato expression (see Supplementary Materials). On day 23, the FosTRAP mice were placed back in context B for the duration of 10 min, following the exact same procedures as during the trigger session, to induce fear memory recall. No shocks were administered during this context re-exposure session. Behavior was videotaped and freezing behavior scored manually by an observer blind to the experimental condition (The Observer XT12, Noldus). Mice were sacrificed 90 min post re-exposure under anesthesia (5% isoflurane inhalation followed by i.p. injection with 200 μl pentobarbital) by perfusion with PBS followed by 4% PFA. The brains were surgically removed and post-fixed for 24 hours in 4% PFA, after which they were transferred to 0.1 M PBS with 0.01% sodium azide and stored at 4 °C.

#### Immunohistochemistry

Right hemispheres of the brains of the PTSD-like (n=9), resilient (n=8), and intermediate (n=17) animals were sliced into 30 μm coronal sections and immunostained for c-Fos and somatostatin protein. Further details are given in the Supplementary Materials.

### Statistical Analyses

Data were analyzed using IBM SPSS Statistics 23. Data points deviating more than three inter-quartile ranges from the median were considered outliers and removed from further analysis (see Supplementary Materials for the exact data points excluded). Normality was checked using the Kolmogorov-Smirnov test. Comparisons between PTSD-like and resilient animals were done using independent sample t-tests when assumption for normal distribution was met. In case the assumption for normal distribution was not met, a Mann-Whitney U test was used to compare PTSD-like vs resilient mice. In case of repeated measures within an animal (i.e., effects of dorsal-ventral axis or distance to soma in case of morphological data) a repeated measures ANOVA was used. In case Mauchly’s test for sphericity indicated that sphericity could not be assumed, Greenhouse-Geisser tests were reported. Results were considered significant if p<0.05. Significant ANOVA group x axis interaction effects were followed up with Bonferroni-corrected *post hoc* t-tests, requiring a p<0.025 to reach significance (p=0.05/2, based on two post hoc tests for the dDG and vDG). Note that uncorrected p-values are reported throughout the text. Figures show average ± standard of the mean (SEM) in case of normally distributed data, and median ± interquartile distances in case of non-normal distribution.

## RESULTS

### Ventral DG spine density is reduced in PTSD-like animals

To assess potential differences in DG neuronal morphology associated with differential susceptibility to PTSD-like symptoms following trauma, a batch of 24 mice was exposed to the PTSD induction protocol and, following a week of recovery, assessed on PTSD-like symptomatology (Fig. S2). DG neuronal morphology of PTSD-like (n=4) and resilient animals (n=5) was assessed by Golgi staining (Fig. 1A). Sholl analyses revealed no significant differences between groups in the total dendritic material in the dorsal and ventral DG (dDG: F(1,7)=1.041, p=0.342, vDG: F(1,7)<1), nor in its distribution across distance to soma (group x distance interaction. dDG: F(24,168)=1.010, p=0.456, vDG: F(22,154)=1.037, p=0.424), whereas there was a clear effect of distance to soma (dDG: F(24,168)=62.784, p<0.001, vDG: F(22,154)=50.742, p<0.001, Fig. 1B). Spine density was not different between PTSD-like and resilient animals in the dDG (U=10, p=1.000), but PTSD-like animals displayed a slightly, but significantly, reduced spine density in the vDG (U=0, p=0.029, Fig. 1C).

**Figure 1.**
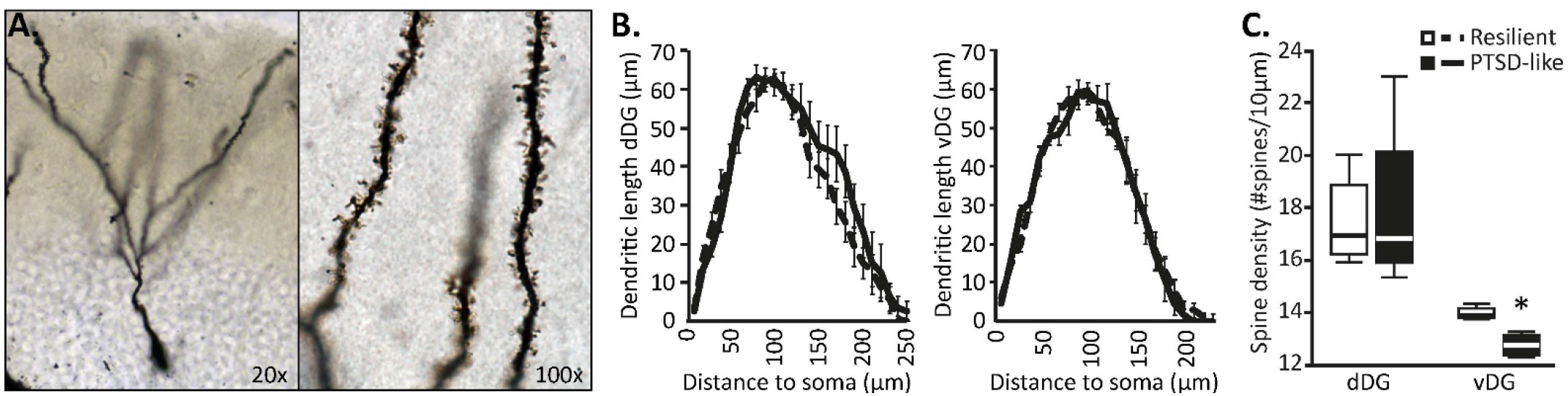
Results for behavioral cohort 1. Reconstruction of Golgi-stained dentate gyrus (DG) granule cells (**A**) revealed no differences in dendritic length between in the dorsal (dDG) or ventral DG (vDG) (**B**), but indicated reduced spine density in the ventral DG of PTSD-like animals, whereas no differential spine density was observed in the dorsal DG (**C**). Behavioral results for this cohort are depicted in Figure S2. *: p<0.05

### *Homer1b/c* expression is reduced in the ventral DG of PTSD-like animals

Next, potential differences in DG gene expression associated with differential susceptibility to PTSD-like symptoms following trauma were assessed in a new batch of 48 mice. To test whether pre-trauma anxiety constitutes a risk factor for later PTSD development, all animals of this batch were additionally tested for anxiety-like behavior prior to trauma exposure.

Animals later categorized as PTSD-like (n=10) vs resilient (n=12) (Fig. S3) did not display different behavior in the open field test prior to trauma exposure. The distance traveled through the center, total distance traveled, the number of crossings through the center, the time spent in the center (all t(20)’s<1) and the latency to enter the center (t(20)=1.581, p=0.129) of the open field (t(20)<1) were not different between groups (Fig. S4A). However, groups did differ in the distance traveled on the open arms of the elevated plus maze (t(19)=2.307, p=0.033), with the PTSD-susceptible animals walking a shorter distance (mean±stdev: 137.56±51.02 cm) compared to the resilient animals (197.99±64.84 cm). No differences were observed in the time spent on the open arms (t(20)=1.395, p=0.178), nor in the total distance traveled on the maze (t(11.812)=1.005, p=0.335, Fig. S4B). Interestingly, significant behavioral differences between PTSD-like vs resilient groups were also observed during the trigger session, where the PTSD-like mice showed a shorter latency to start freezing than resilient animals (U=22, p=0.017). Furthermore, a majority of PTSD-like mice –in contrast to resilient mice– showed freezing behavior before the first foot shock was administered in this novel context (U=88, p=0.020, Fig. 2A). Subsequent shock-induced freezing was not different between PTSD-like and resilient ones (t(20)=1.216, p=0.238, Fig. 2A), suggesting similar threat coping mechanisms.

**Figure 2.**
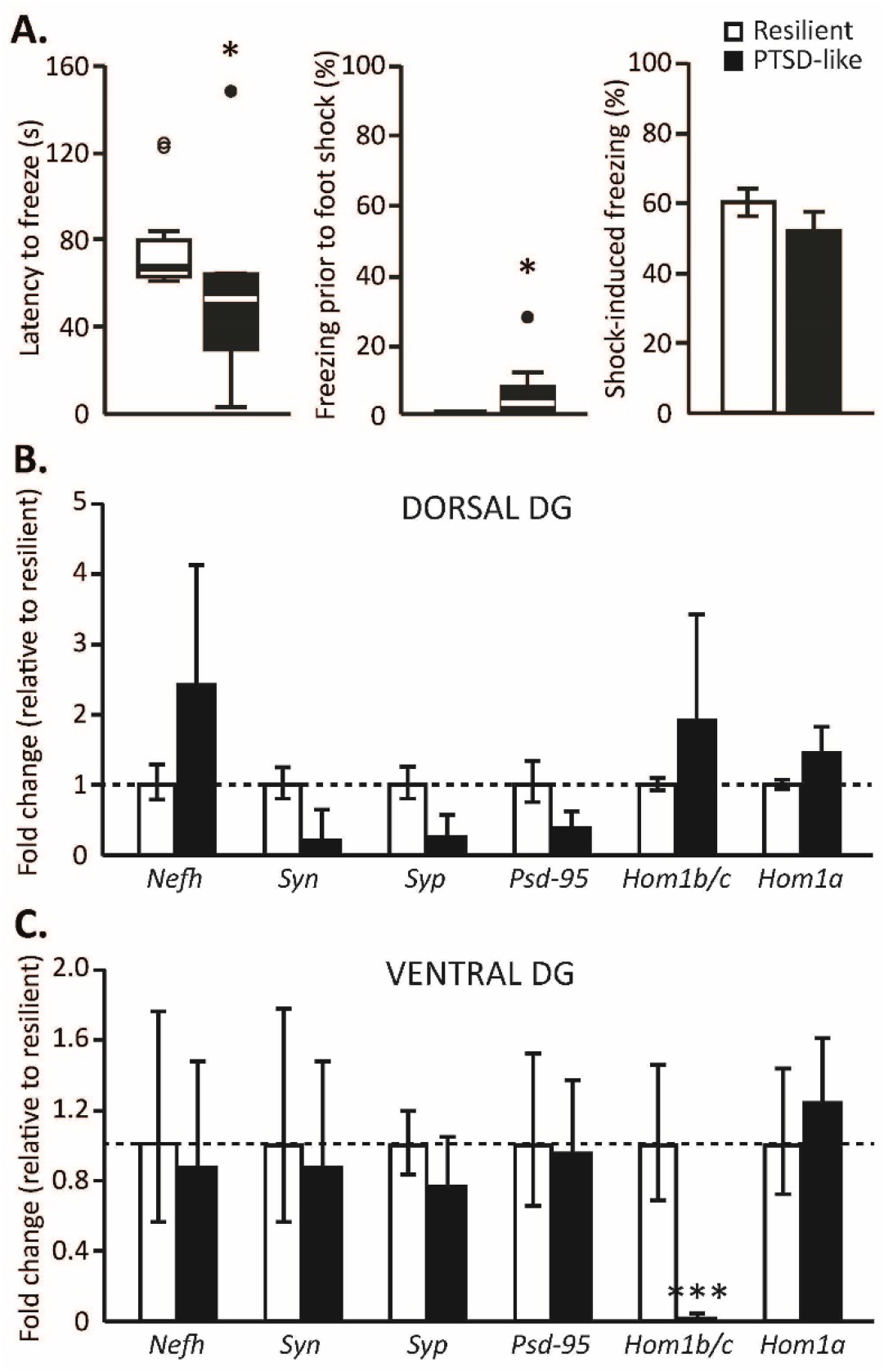
Results for behavior cohort 2. PTSD-like animals showed a lower latency to freeze in the trigger context, with substantial freezing already prior to the first foot shock in this novel context. Subsequent shock-induced freezing was not different between groups (**A**). Gene expression levels of synaptic proteins were not different between PTSD-like and resilient animals in the dorsal dentate gyrus (DG) (**B**), but revealed a strong reduction in the expression of the Homer1b/c gene (*Hom1b/c*) in the ventral DG (**C**). Behavioral results for this cohort on PTSD symptoms are depicted in Figure S3, whereas anxiety measures are displayed in Figure S4. *Nefh*: neurofilament H, *Syn*: synapsin I, *Syp*: synaptophysin, *Psd-95*: postsynaptic density-95, *Hom1a*: Homer1a splice variant, ***: p<0.001

We continued with analyzing the DG’s expression levels of genes encoding excitatory presynaptic (*Syn* and *Syp*) and postsynaptic (*Homber 1b/c* and *Psd-95*) markers related to spine density, a spine-localized immediate early gene (*Homer 1a*) and a neuronal marker (*Nefh*), previously found to be affected by stress^35,45^. The dorsal (Fig. 2B) and ventral DG (Fig. 2C) did not reveal differential expression of *Nefh* mRNA between PTSD-like vs resilient mice (p’s>0.13), suggesting that the number of neurons did not differ between groups^35^. Moreover, no differences were found in mRNA levels of the presynaptic vesicle markers *Syn* and *Syp* (p’s>0.1), indicating that synaptic release in the PTSD-like mice was not altered. Interestingly, mRNA levels for *Homer1b/c* showed a main effect of axis (F(1,13.058)=41.810, p<0.001), group (F(1,14.981)=4.868, p=0.043) and a group x axis interaction (F(1,13.058)=34.321, p<0.001), which was caused by substantially lower expression in the vDG of PTSD-like animals compared to resilient animals (t(7.388)=3.475, p=0.009), but not the dDG (t(6.285)=1.107, p=0.309). These effects seemed to be driven by 3 PTSD-like animals that showed extremely low levels of Homer1b/c expression (<1% of the levels of resilient animals), but exclusion of these animals still revealed significant group differences (t(13)=2.540, p=0.025). No differences in the *Homer1a* splice variant were observed between groups (p’s>0.35). Moreover, no group differences in *Psd-95* mRNA levels were observed (p’s>0.23).

### PTSD-like animals activate a larger DG neuronal ensemble during trauma encoding

Next, we wanted to investigate whether differential trauma susceptibility was also related to distinct DG functionality in terms of its activity during trauma memory encoding and retrieval. To do so, we used FosTRAP mice^36^, in which the injection of tamoxifen induces the expression of the fluorescent marker tdTomato in all c-Fos-expressing (i.e., activated) neurons. For the study of DG function, the FosTRAP mouse line was preferred over the ArcTRAP mouse line^36,46,47^, as the latter is characterized by substantial background staining of DG neurons in non-injected animals (i.e., labeled neurons in the absence of tamoxifen (Fig. S5)), which is not observed in FosTRAP mice (Fig. S6)^36^. Pilot experiments indicated that FosTRAP mice showed no alterations in fear behavior or memory performance, and significant labeling of DG neuronal activity upon tamoxifen injection (Fig. S6, S7), qualifying them for the experiment. To investigate whether DG activity during trauma encoding can predict PTSD susceptibility, we injected 40 FosTRAP mice with tamoxifen prior to trauma induction, followed by the trigger the next day. Again, following a week of recovery, mice were tested for PTSD-like symptoms, dissociating susceptible (n=9) from resilient (n=8) mice (Fig. S8). To additionally test whether PTSD susceptibility is associated with altered DG recruitment during the recall of the traumatic experience, animals were re-exposed to the trigger context prior to sacrifice, and their brains analyzed for recall-induced c-Fos expression.

Retrospective analyses revealed no differential locomotor behavior in the trauma context between PTSD-like vs resilient mice (overall mobility; t(12)<1). However, significant differences were again observed between groups during the trigger session, when the PTSD-like mice showed a shorter latency to start a freezing bout (defined as a period of complete immobility for >2s) than resilient animals (t(8.910)=2.374, p=0.042), and on average started freezing well before the first foot shock administration in this novel context (Fig. 3A). Overall freezing prior to the first shock administration was not different between PTSD-like and resilient animals (t(13)<1), nor were subsequent shock-induced freezing levels (t(12.786)<1, p=0.561, Fig. 3A). Following the behavioral categorization, mice were re-exposed to the trigger context to induce fear memory recall. In contrast to the shorter latency to start freezing observed during the trigger exposure, PTSD-like animals tended to show a somewhat longer latency to start freezing during its recall (t(8.314)=2.119, p=0.066, Fig. 3B). Examination of the freezing levels of PTSD-like and resilient animals over time revealed a significant effect of time (F(3.897,54.561)=3.081, p=0.024), with freezing reducing upon prolonged context exposure, but no differences between groups (main effect of group; F(1,14)=1.279, p=0.277, group x time interaction; F(3.897,54.561)<1, Fig. 3B).

**Figure 3.**
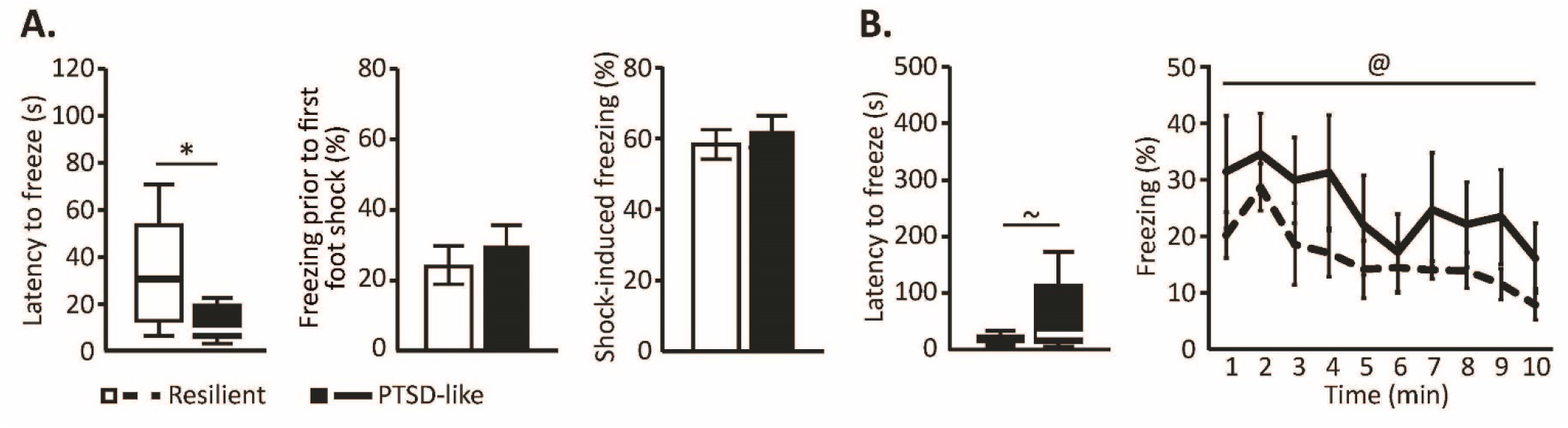
Behavioral assessment of cohort 3 confirmed a shorter latency to freeze in PTSD-like animals in the trigger context (**A**), and revealed a trend towards a longer latency to freeze upon re-exposure to this context compared to resilient animals (**B**), with no differences in overall freezing levels or their reduction across prolonged exposure between these groups. Behavioral results on PTSD symptoms for this cohort are depicted in Figure S8. *: p<0.05, ∼: p=0.066, effect of PTSD-phenotype; @: p<0.05, effect of time

Analysis of the DG neuronal populations active during the trauma and trigger exposure (i.e., the amount of tdTomato-expressing neurons) in PTSD-like vs resilient mice revealed that the PTSD-like animals displayed a significantly larger active neuronal ensemble during the PTSD induction protocol compared to resilient animals (main effect of group; F(1,12)=4.841, p=0.048), and that this difference was independent of ventral-dorsal axis (group x axis interaction; F(1,12)<1, Fig. 4B). The amount of active DG neurons during fear memory recall as assessed by c-Fos expression was not significantly different between groups (main effect of group; F(1,12)<1, group x axis interaction; F(1,12)<1, Fig. 4C). In terms of the reactivation rates (defined as the number of neurons double-positive for tdTomato and c-Fos divided by the total number of tdTomato positive cells^48,49^), no significant group differences were observed for the dDG (median ± interquartile range: dDG_resilient_=0.44±2.20%, dDG_PTSD-like_=0.00±1.21%, U=19.5, p=0.573), whereas the vDG revealed trend-level significant higher reactivation in PTSD-like vs resilient mice (median ± interquartile range: vDG_resilient_=0.00±0.00%, vDG_PTSD-like_=0.41±1.88, U=32.5, p=0.065). However, reactivation rates were overall very low. Remarkably, the number of DG somatostatin neurons –primarily located in the DG hilar region (Fig. 4A)– was significantly lower in the dDG of PTSD-like vs resilient mice (t(12)=2.691, p=0.020), whereas a trend towards increased counts of somatostatin neurons was observed in the vDG (t(11)=1.845, p=0.087), resulting in a significant group x axis interaction (F(1,10)=8.511, p=0.015, Fig. 4D). Both the number of active DG somatostatin neurons during encoding and remote recall were very low (Table S1) and not different between groups (all p’s>0.5).

**Figure 4.**
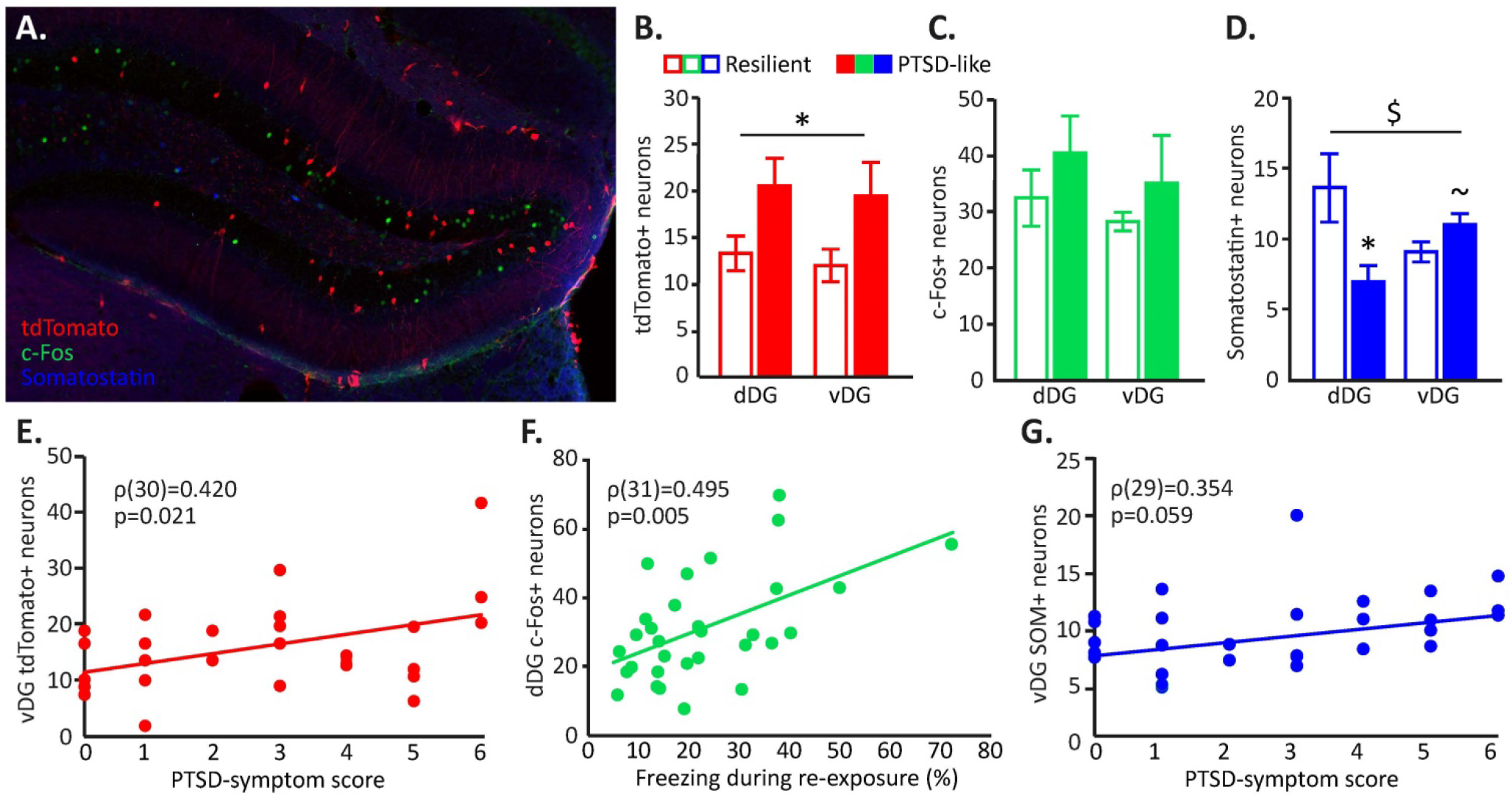
In cohort 3, dentate gyrus (DG) activity during trauma memory encoding (marked by tdTomato expression), remote trauma memory recall (marked by c-Fos expression), as well as somatostatin (SOM) interneuron levels were assessed by immunohistochemistry (**A**). PTSD-like animals displayed an increased population of DG neurons active during trauma encoding (**B**), but no differences during its remote retrieval (**C**). The number of somatostatin neurons identified in the dorsal (dDG) and ventral (vDG) differed between groups as well (**D**). Correlational analyses involving also mice with an intermediate PTSD-phenotype revealed associations between specifically the ventral DG and PTSD symptom score (**E**,**G**), whereas the dorsal DG seemed to relate to trauma memory strength (**F**). Quality checks for the TRAP method are presented in Figures S5 to S7. *: p<0.05, ∼: p=0.087, effect of PTSD-phenotype; $: p<0.05, hippocampal axis x PTSD-phenotype interaction

Based on significant correlations between overall PTSD symptom score and the number of active neurons in the vDG during the trauma and trigger exposure (i.e., number of tdTomato positive cells; ρ(14)=0.703, p=0.005), as well as the number of somatostatin neurons within the vDG (ρ(13)=0.623, p=0.023), we additionally analyzed the brains of 17 animals that showed an intermediate PTSD phenotype (1≤PTSD symptom score≤4). All DG outcome measures were subsequently tested for significant associations with PTSD symptom score, instead of group differences between the extreme scoring animals only. These analyses confirmed previous findings of a larger active DG population during trauma+trigger processing predicting greater PTSD-like symptomatology (ρ(30)=0.426, p=0.019), and suggested that this correlation was particularly prominent for the vDG (ρ(30)=0.420, p=0.021, Fig. 4E), whereas the association between PTSD symptom score and dDG activity failed to reach significance (p=0.153, Fig. S9A). Furthermore, these analyses confirmed an increased presence of somatostatin neurons in the vDG (but no differences in the dDG; ρ(29)=-0.028, p=0.884, Fig. S9B) in the development of PTSD symptomatology, although this association reached trend-level significance only (ρ(29)=0.354, p=0.059, Fig. 4G). When testing for correlations between the number of active DG neurons and trauma memory recall, DG activity during trauma encoding appeared not predictive of later re-exposure induced freezing (p’s>0.5), but increased recall-induced activity in the dDG (ρ(31)=0.495, p=0.005, Fig. 4F) (but not vDG (p=0.236)) was associated with enhanced memory recall. These data implicate particularly the dDG in fear memory recall, which seems unaffected in PTSD-like compared to resilient animals. Deviations in vDG function however seem to correlate more strongly with differences in PTSD symptomatology.

## DISCUSSION

We investigated the association between DG structure and function and the susceptibility to develop PTSD-like symptoms following trauma. Besides trauma-induced PTSD-like symptomatology, PTSD-like animals seemed to displayed some increased anxiety-like behavior already prior to trauma, and greater anxiety in a novel context following trauma exposure. No clear differences between phenotypes were observed in remote trauma memory recall. Comparison of the vDG of PTSD-like vs resilient mice revealed lower spine density, reduced expression of the postsynaptic protein homer 1b/c gene, increased population of neurons active during trauma encoding and increased presence of somatostatin neurons to be associated with PTSD susceptibility. In contrast, the dDG of PTSD-like animals did not differ in terms of spine density or synaptic protein gene expression, but displayed more active neurons during trauma encoding and less somatostatin neurons. As such, these data seem to implicate mainly the vDG in establishing PTSD-like symptoms of trauma-related arousal in this animal model.

Elucidating the neurobiological basis of individual differences in PTSD susceptibility has been a central question in stress-related research. Here, we used an animal model to investigate this, showing substantial heterogeneity in the behavioral consequences of trauma exposure (i.e., risk assessment, anxiety, hypervigilance, pre-pulse inhibition and activity during the inactive phase) in mice. Noteworthy, mice were classified as PTSD-like or resilient based on a compound score comprising multiple behavioral PTSD-like symptoms, rather than single behavioral features. This classification resembles the situation in patients^50^, which can be diagnosed with PTSD based on 20 criteria across four distinct symptom categories, resulting in a highly heterogeneous patient population (DSM-V^1^). Accordingly, we observed substantial behavioral variability both within and across the three different cohorts in this study, as well as when comparing our findings to previous reports on this PTSD model^39,40^. However, altered vDG function/structure was observed for all three cohorts, supporting its involvement in a wide range of PTSD symptoms.

Dissociating PTSD-like vs resilient animals, we found susceptible mice traveled shorter distance on the open arms of the elevated plus maze, indicative of a reluctance to explore relative danger zones; indicative of increased anxiety. However, no behavioral differences were observed in the open field, a potentially less threatening environment. These findings match prior animal work on reduced exploratory drive^51^ and enhanced anxiety^52^ predicting trauma sensitivity, as well as human reports on trait anxiety being predictive of PTSD risk and symptom severity^53,54^. Furthermore, PTSD-like animals displayed increased behavioral freezing upon exposure to the unfamiliar trigger context post-trauma. This is in line with reports on elevated distress/arousal soon after trauma being predictive of later PTSD symptom severity^55,56^ and intrusions^57^. It is tempting to relate the increased novelty-induced anxiety to generalized fear in PTSD-susceptible mice, as has been reported by others^58^, but future dedicated assays on the extent to which fear generalizes across contexts is required to warrant such a claim.

PTSD-like animals revealed decreased spine density specifically in the vDG, but no differences in dorsal or ventral DG dendritic length. Alterations in DG morphology have been linked to inter-individual differences in stress susceptibility before, with only the animals most susceptible to trauma^32,34^, learned helplessness^33^, or chronic social defeat^59^ showing reductions in DG spine density. In our PTSD mouse model these effects seem to apply to the vDG specifically. Similarly, we only observed a reduction in the expression of the postsynaptic protein homer 1b/c gene in the vDG. Homer 1b/c is an excitatory postsynaptic density scaffolding protein^44^, regulating spine morphogenesis, synaptic plasticity and the stabilization of synaptic changes during long-term potentiation (LTP)^60^; suggesting an active role in behavioral plasticity^61^. Its hippocampal expression levels have been found reduced following traumatic stress, and associated with generalized fear^35^. Previous work has also described alterations in the other synaptic genes assessed following stress or trauma^35,45^, but our study is the first in assessing alterations in specifically the DG and contrasting susceptible vs resilient individuals, providing more nuance to earlier findings. Overall, our assessments of DG synaptic contacts suggest altered vDG synaptic signaling of glutamatergic input, without any dDG differences.

Importantly, larger active populations of DG neurons during trauma encoding predicted later PTSD symptoms, predominantly in the vDG, where the number of active neurons significantly correlated with PTSD symptom score. These findings correspond with the suggested role for the vDG in mediating anxiety-related behaviors^18,21-23^, and a reduction of vDG neuronal activity during anxiogenic situations conferring stress resilience^62^. In contrast, larger dDG neuronal populations active during trauma encoding predicted increased freezing upon remote memory recall, which supports its critical role in fear memory acquisition^18,63,64^. DG activity is under tight control of local GABAergic interneurons^11,15,65^, with a prominent role for somatostatin-expressing interneurons^16^, which increase the threshold of input required for acquisition of new memories, filtering out irrelevant environmental cues^66^. Correspondingly, activation of somatostatin neurons has been found to reduce the size of the activated granule cell population upon encoding to ensure memory specificity^16^. The reduction in dDG somatostatin neurons and increased number of DG granule cells fit these observations. No differences between phenotypes were observed in DG activity during memory recall, nor reactivation rates. Although reactivation of DG neurons activated during encoding has been shown to suffice to induce recent memory recall^67,68^, memories are known to reallocate to either cortical representations over time^69,70^, or to different cells within the hippocampus^71^. This might explain our low DG reactivation rates upon remote recall^46^. Altogether, these findings suggest that aberrant activity of the vDG is implicated in establishing PTSD trauma-related and arousal symptoms modeled in our mouse model, whereas the dDG seems to be more involved in actual trauma memory processing, which seems unaffected here. Future studies testing for fear generalization/pattern separation upon re-exposure to a broad array of contexts, as well as the capacity for fear extinction –in which the DG is also involved^72,73^– should further determine whether trauma memory processing is affected as well.

Findings of an increased vDG population of active neurons during trauma memory processing may seem at odds with the increased potential for inhibition (i.e., more somatostatin neurons) and reduced capacity for excitation (a reduction in glutamatergic spines and excitatory postsynaptic scaffolding protein gene expression). Noteworthy, all assessments of DG excitatory/inhibitory structural markers have been obtained post-trauma, posing the question on whether these are cause or consequence of the acquired symptomatology. Previous research has implicated aberrant hippocampal function and structure as both^74,75^. Therefore, it could well be that the observed alterations in excitatory/inhibitory regulation reflect a compensatory response to an initial excess of excitatory input^76^. However, prior observations that particularly vDG granule cell morphology is related to overall anxiety-like behavior, independent of an animal’s stress history^31^, may suggest that these alterations are rather cause than consequence of trauma-associated symptoms. Interestingly, we also observed a significant correlation between vDG homer 1b/c gene expression and pre-trauma anxiety (distance moved on open arm; ρ(17)=-0.498, p=0.042), supporting that the synaptic differences are a pre-disposing trait rather than a state marker of pathology.

Some limitations should be mentioned. Firstly, the structural assessments of glutamatergic/GABAergic regulation cannot directly be related to the neurons involved in trauma memory processing, as these implicate generic DG changes independent of the functional population. Future studies should further investigate this by analyzing morphology and gene expression patterns of trauma-activated neurons specifically. Secondly, we did not consider the heterogeneity of DG granule cells in terms of age, whereas particularly newborn neurons seem to be involved in pattern separation^77,78^. Thirdly, similarly to other hippocampal memory engram labeling studies^67,79,80^, almost exclusively excitatory cells were labeled by tdTomato expression, leaving the contribution of local interneurons unresolved. Lastly, further manipulation studies will be necessary to elucidate a causal link between the observed vDG alterations and trauma-related behavior.

Concluding, we found little evidence for aberrant dDG structure and function being related to PTSD-like symptomatology. In contrast, the vDG displayed several deviations indicative of aberrant glutamatergic and GABAergic regulation of granule cell activity in PTSD susceptible mice compared to those that are resilient. These changes appeared associated with elevated anxiety-like behavior even prior to trauma exposure, and higher (generalized) fear to novel contexts. Thereby, the vDG seems critically involved in establishing the PTSD symptoms as assessed in our mouse model, and an important target for further research into the psychopathology of PTSD.

## Acknowledgements

M.J.A.G.H. was supported by Veni Grant 863.15.008 and J.R.H. by Vidi Grant 864.10.003 awarded by the Netherlands Organization for Scientific Research.

