## Supplementary Materials for "Aberrant ventral dentate gyrus structure and function in individuals susceptible to post-traumatic stress disorder"

*PTSD Model.*

*Trauma and trigger events.* Shock exposure was conducted in fear-conditioning boxes (TSE Systems, Bad Homburg, Germany) for experiments 1-2, whereas in experiment 3 operant touch screen chambers connected to a shock generator (Campden Instruments) were used. The latter chambers were equipped with infrared beams located on both sides of the chambers. Beam break data were analyzed and used as a proxy for locomotor activity during the trauma session. For all experiments, to generate the specific context A, mice were moved to a dark experimental room per two/three in dark, carton boxes and placed individually in the shock boxes. As context, dark, triangular shaped inserts were placed in the chamber with a steel grid and metal tray at the bottom. Chambers were sprayed with 1% acetic acid, and a continuous 70 dB background noise was present during the entire session. For context B, mice were moved per 2/3 in clean home cages to the 70 lux illuminated experimental room and placed individually in the shock boxes, which were contextually altered containing a curved white insert and a white tray underneath the metal grid. Chambers were sprayed with 70% ethanol and during the session the house light in the box was turned on without additional background noise.

*Behavioral testing.*

*Dark-light transfer test*. On day 8 of the protocol, mice were tested in the dark-light transfer test[40]. The test was executed in a box that was divided into a dark compartment (29 x 14 cm) and brightly illuminated (~1 000 lux) compartment (29 x 29 cm), which were connected by a retractable door. The mice were placed in the dark compartment of the box, the door was opened and the 5 min test session started. Movements of the mice were recorded and scored automatically using Ethovision XT (Noldus). The time spent in the risk assessment area, a small area by the opening of the door of the light compartment (6 x 3 cm), was measured to calculate the percentage risk assessment; the amount of time spent in the risk assessment zone as a percentage of total time spent in the lit arena outside of that zone. *Marble burying.* At day 10, mice were placed in a 5-10 lux illuminated black open box (30 x 28 cm), filled with a 5 cm layer of corncobs and 20 marbles centrally arranged (4 x 5) on top of that layer. Mouse burying behavior was videotaped for 25 min. Videos were scored by assessing the amount of buried marbles after 25 minutes.

*Acoustic Startle and Pre-pulse Inhibition (PPI).* At day 12, mice were individually placed in small, transparent constrainers, mounted on a vibration sensitive-platform, which contained a high-precision sensor for detecting movement, inside a ventilated cabinet that contained two high-frequency loudspeakers (SR-LAB, San Diego Instruments). Movements of the mice were measured with a sensor inside of the platform. The test started with an acclimatization period of 5 min in which a background noise of 70 dB was presented, which was maintained throughout the session. Thirty-two startle responses of 120 dB, 40 ms in duration and with a random varying ITI (12-30 s) were presented with another 36 startle responses preceded 100 ms earlier by a 20 ms pre-pulse of either 75 dB, 80 dB or 85 dB. The latency to peak startle amplitude was assessed (i.e., time to Vmax) together with the percentage of PPI; [1 - (mean pre-pulse startle amplitude / mean startle amplitude without pre-pulse) x 100].

*Locomotion during light phase*. Immediately after the pre-pulse inhibition test, mice were single housed in Phenotyper cages (45 x 45 cm, Noldus) for 72 hours while their locomotion was being recorded by an infrared-based automated system (EthoVision XT, Noldus). The first 24 hours was considered as habituation time and data were discarded. Total locomotion during the two last light phases was assessed.

*Experiment 1: DG Neuronal morphology.*

*Neuronal Reconstruction and Spine Density.* A Zeiss Axioskop FS microscope (Zeiss, Oberkochen, Germany) with an 100X oil-immersion objective lens (plan-NEOFLUAR, Zeiss, Oberkochen, Germany) and the program Neurolucida (MBF Bioscience, Microbrightfield Inc., Williston, VT, USA) were used by an observer blind to the experimental conditions to examine the neuronal morphology of dorsal (Bregma -1.46:-2.18 mm) and ventral (Bregma -2.92:-3.40 mm) DG granule cells. For every animal, 5 cells per region were selected applying the following criteria: must come from the target region within the specified distance from Bregma, must be completely impregnated, must contain a clear soma and primary dendritic branch distinguishable from surrounding neurons. The Neurolucida software was used to trace the entire apical dendritic tree of the selected cells, after which spines were counted in an average of 6 segments approximately 50 μm in length at variable branch order (range: 3-11) and distance to soma (range: 20-360 μm). No distinction was made between different spine morphologies. Average number of spines per 10 μm was calculated for all segments and averaged to obtain a single spine density measure per animal per hippocampal subregion. It should be noted that the spine density values obtained are likely to be an underestimation of the actual density because spines protruding beneath or above the dendritic segment were not visible and thus not accounted for in this analysis.

*Experiment 2: DG Gene Expression.*

*Open field.* To assess pre-trauma levels of anxiety, mice were first tested in the open field test. The open field apparatus consisted of a white Plexiglas box (50 x 50 x 40 cm) lightened with 120 lux. Each mouse was placed in the corner of the apparatus to initiate a 10 min test session. Time spent in the center (the inner 25 x 25 cm), distance traveled in the center, and total distance traveled were captured using a camera mounted above the apparatus and analyzed by Ethovision software (Noldus, Wageningen, Netherlands).

*Elevated plus maze.* As a second test for pre-trauma anxiety, the slightly more aversive elevated plus maze was used. The elevated plus maze comprised a central part (5 x 5 cm), two opposing open arms (30.5 x 5 cm), and two opposing Plexiglas closed arms (30.5 x 5 x 15 cm), elevated at a height of 53.5 cm. The open arms were illuminated with 6-9 lux. Mice were placed in one of the closed arms facing the center to initiate a 5 min test session. Time spent in the open arms, distance traveled in the open arms, and total distance traveled were captured using a camera mounted above the apparatus and analyzed by Ethovision software (Noldus, Wageningen, Netherlands).

*Quantitative PCR.* Forward and reverse primers were specifically designed for the target genes using the UCSC genome browser (http://genome-euro.ucsc.edu) and primer3 (http://frodo.wi.mit.edu/primer3). Multiple exons were included in the primer design to afford more stability and specificity of product. *Syn1*: CCACAGAGAATCCACCATTG (forward), ATCGACGAACCGCACAC (reverse); *Syp*: GTGGAGTGTGCCAACAAGAC (forward), TTGGTAGTGCCCCCTTTAAC (reverse); *Psd-95*: CCGCTACCAAGATGAAGACAC (forward), ATATCCTGGGGCTTCTAGGG (reverse); Homer1a: CCAC-AGGCCTACAGAGTTTTC (forward),

CTGCCATCAATGCTAACAGG (reverse); Homer1b/c: ATGAGAGAA-CACCCGATGTG (forward), GAGAGAGGGCCTGGGTTC (reverse); *Nefh*: TCCCACTTGGTGTTTCTCAG (for-ward), AGGCAGACATTGCCTCCTAC (reverse); *Hprt*: TGTTGTTGGATATGCCCTTG (forward), CAACTTGC-GCTCATCTTAGG (reverse); *Cyc1*: CCTTCATTCTGCCGTGAGTG (forward), CACACCTGCTTGTATACCTGG (reverse). qPCR was performed using the SensiFAST™ SYBR® No-ROX Kit (Bioline, Taunton, MA, USA). Master mixes and cDNA dilutions were prepared by hand and reaction mixes were compiled with QIAgility (QIAGEN, Venlo, the Netherlands) and run in a Rotor-Gene Q with Rotor-Disc 100 (QIAGEN, Venlo, the Netherlands). Because of the SYBR, the green channel was used. The protocol for the qPCR runs included 40 cycles of alternating 20 s at 64 °C and 5 s at 95 °C. Melting curve analysis was included to verify the specificity of primer binding. Every individual reaction contained 5 µL kit, 0.4 µL each of forward and reverse primer solution (10 µM), 2.2 µL RNase-free water and 2 µL cDNA solution (0.1 ng/µL) for a total reaction volume of 10 µL.

*Experiment 3: DG Neuronal Activity.*

*Tamoxifen*. FosTRAP mice were injected with tamoxifen solution 7 hours prior to trauma exposure. Tamoxifen was dissolved in a 10% ethanol/corn oil solution at a concentration of 10 mg/mL by overnight sonication and stored at -20°C until further use. At the day of injection, solutions were heated to body temperature and injected intraperitoneally at a dosage of 150 mg/kg.

*Immunohistochemistry.* Right hemispheres of the brains of the PTSD-like (n=9), resilient (n=8), and intermediate (n=17) animals were first cryoprotected in 30% sucrose solution and subsequently sliced at 30 µm thickness using a freezing sliding microtome (Microtom HM440E, GMI Inc., Ramsey, MN, USA) and stored in 1x PBS with 0.01% sodium azide. Free floating immunohistochemistry was performed on 4-6 sections of both the dorsal DG (Bregma -1.46:-1.94 mm) and ventral DG (Bregma -2.92:-3.52 mm). Sections were washed three times in 1x PBS and blocked in PBS-BT (1x PBS with 0.3% Triton X-100 and 1% bovine serum albumin (BSA)) for 30 minutes at room temperature (RT). Incubation of the primary antibody was performed overnight (guinea pig anti-c-fos, 1:750, 226004, Synaptic Systems; rat anti-somatostatin, 1:200, MAB354, Merck Chemicals) in PBS-BT for 18 hours at room temperature (RT). Sections were washed three times in 1x PBS, and incubated with the secondary antibody (Alexa 647-conjugated donkey anti-guinea pig, 1:400, AP193SA6, Merck Chemicals; Alexa 488-conjugated donkey anti-rat, 1:200, A-21208, Thermo Fisher) for 3 hours at RT. Lastly, slices were washed three times in 1x PBS, mounted on gelatin-coated slides using FluorSave$^{TM}$ reagent (Merck Chemicals) and cover slipped.

*Image acquisition and cell counting.* For cell counting, images of at least 4 sections of each DG region were captured through a light microscope (Axio Imager 2, Zeiss) using a 10x objective lens and a LED module (Colibri 2, Zeiss). Cells were manually counted per region in Fiji software by an experimenter blind to the experimental group. DG region size/length was assessed and corrected for to obtain standardized measures of cell density. Normalized cell counts were averaged per region per animal and subjected to statistical testing.

*Statistical analysis.*

Data points were considered outliers when deviating >3 inter-quartile ranges from the median, and were therefore removed from the statistical analyses. This made that for experiment 1, no data points were excluded. For experiment 2, data points on distance travelled on the open arm of the EPM ($n_{PTSD\text{-}like}$=1), total distance travelled on the EPM ($n_{PTSD\text{-}like}$=1), latency to freeze in the trigger context ($n_{resilient}$=1), and latency to freeze upon trigger context re-exposure ($n_{resilient}$=1) were removed. For experiment 3, data points on latency to freeze in the trigger context ($n_{PTSD\text{-}like}$=1), latency to freeze upon trigger context re-exposure ($n_{resilient}$=1), and number of SOM neurons in the dDG ($n_{PTSD\text{-}like}$=1) and vDG ($n_{PTSD\text{-}like}$=1) were removed.

### SUPPLEMENTARY DATA

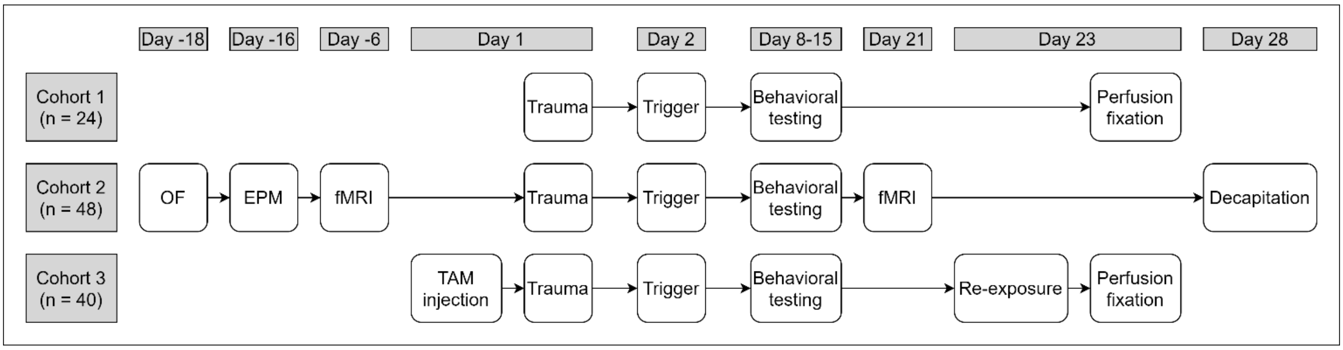


**Figure S1.** Timeline of the behavioral protocols as implemented for experiments 1-3.

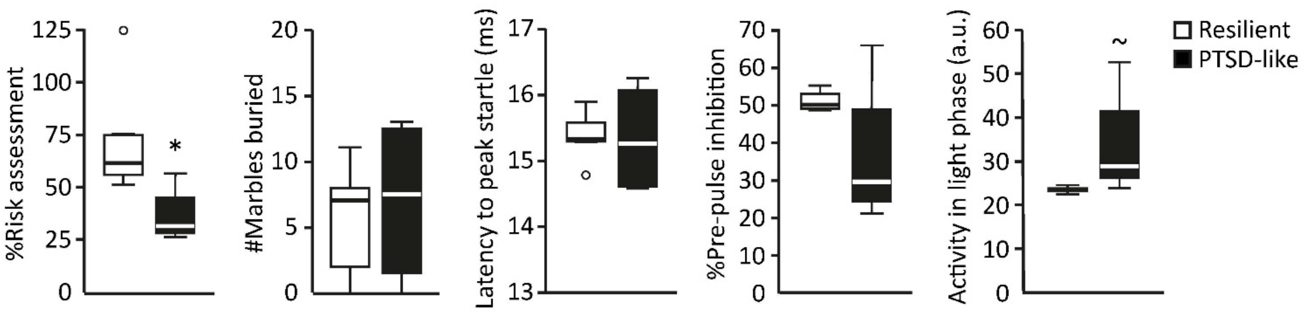


**Figure S2.** Behavioral assessment of the Golgi cohort (experiment 1). PTSD-like animals (n=4) displayed significantly reduced risk assessment behavior (U=2, p=0.050), and a tendency towards increased locomotor activity during the light phase (U=15, p=0.057) compared to resilient animals (n=5). Latency to peak startle (U=10, p=1.000), marble burying behavior (U=12.5, p=0.539), and percentage pre-pulse inhibition (U=4, p=0.343) were however not significantly different between groups. *: p<0.05, ~: p=0.057

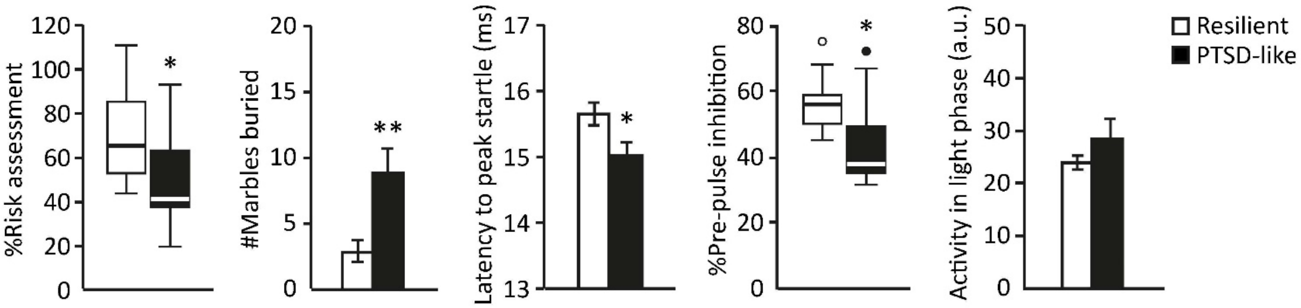


**Figure S3.** Behavioral assessment of the gene expression cohort (experiment 2). PTSD-like animals displayed significantly reduced risk assessment behavior (U=22, p=0.011), increased marble burying (t(13.145)=3.198, p=0.007), shorter latencies to peak startle (t(20)=2.519, p=0.020), and impaired pre-pulse inhibition (U=24, p=0.017) compared to resilient animals. Locomotor activity during the light phase did however not significantly differ between groups (t(20)=1.280, p=0.215). *: $p<0.05$

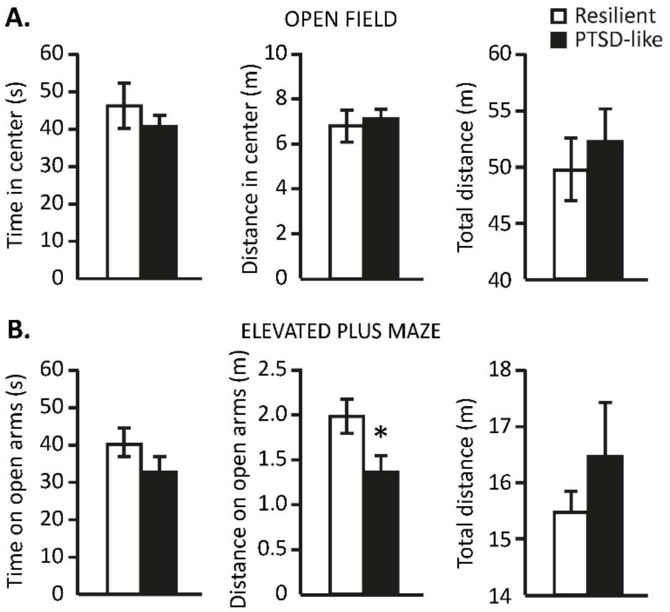


**Figure S4.** Anxiety-like behavior assessed prior to trauma exposure in the open field test (**A**) and elevated plus maze **(B)** revealed that mice developing PTSD-like symptomatology following trauma exposure travelled reduced distance on the open arms of the elevated plus maze. *: $p<0.05$

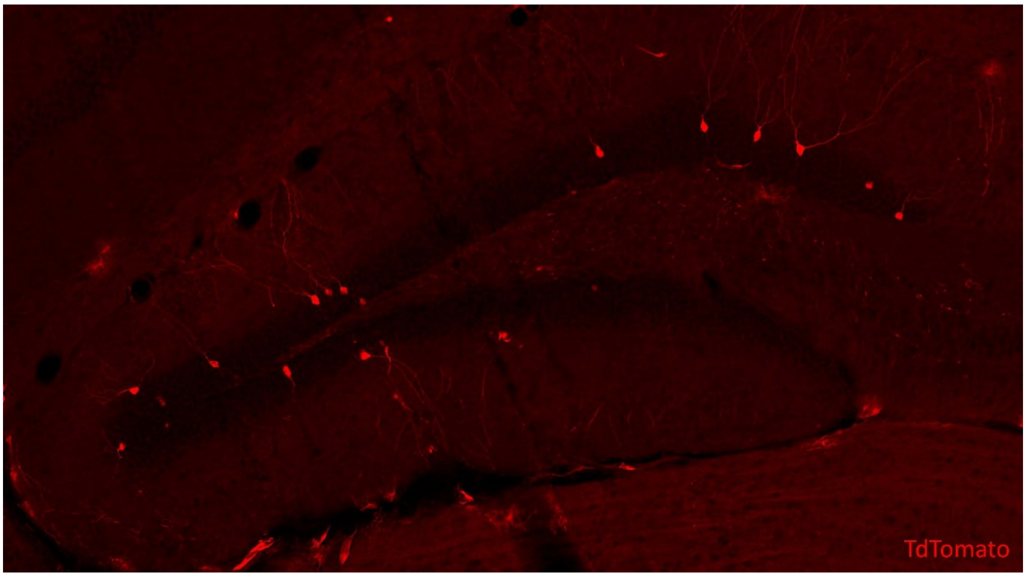


**Figure S5.** ArcTRAP mice displayed tdTomato-labeled neurons in the dentate gyrus even in the absence of tamoxifen.

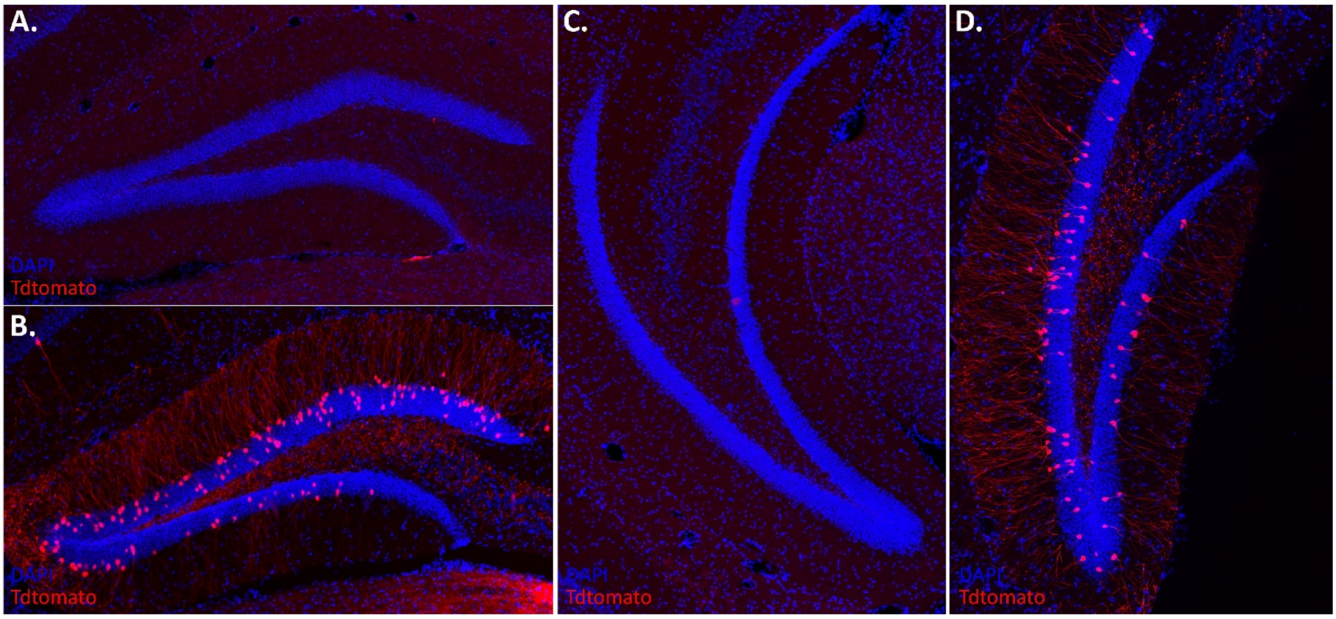


**Figure S6.** FosTRAP mice displayed no neurons labeled by tdTomato expression in the dorsal (**A**) and ventral (**C**) dentate gyrus in the absence of tamoxifen, but clear labeling upon trauma exposure combined with tamoxifen injection (**B**, **D**).

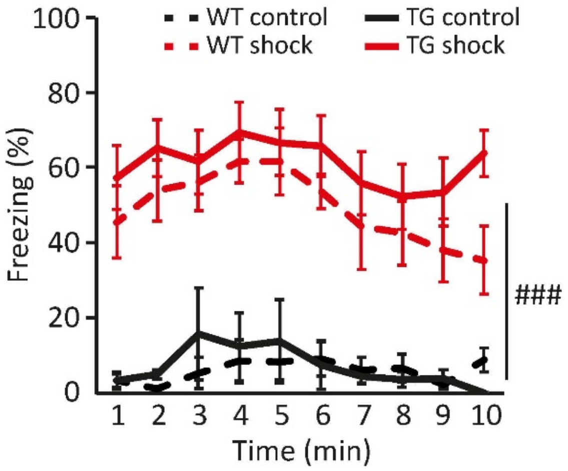


**Figure S7.** To validate that the FosTRAP mice did not show any memory phenotype, as well as to exclude a potential effect of tamoxifen injection on fear learning and memory, 8 FosTRAP mice were injected with tamoxifen and then exposed them to electric foot shocks in a certain context (TG shock). An additional 8 FosTRAP mice were exposed to the same context, but did not receive any shocks (TG no shock). The memory for this episode was tested 5 days later by re-exposing the mice to the same context and the freezing levels of both groups were assessed and compared to their respective non-injected WT-littermates (WT control = 7, WT shock = 7). As expected, mice that previously received foot shocks displayed higher freezing levels than non-shocked animals (###: $F(1,26)=84.754$, $p<0.001$). Moreover, a main effect of time ($F(4.678,121.624)=4.087$, $p=0.002$) was observed without a time x group interaction ($F(4.678,121.624)=1.075$, $p=0.346$), which was best modeled with a quadratic function ($F(1,26)=12.974$, $p=0.001$). Importantly, no effects of genotype were observed on freezing levels (main effect; $F(1,26)=1.475$, $p=0.235$, genotype x group; $F(1,26)=1.050$, $p=0.315$, genotype x time; $F(4.678,121.624)<1$, genotype x group x time; $F(4.678,121.624)=1.351$, $p=0.250$), suggesting that fear memory (and expression) is not influenced by the genotype nor tamoxifen injection.

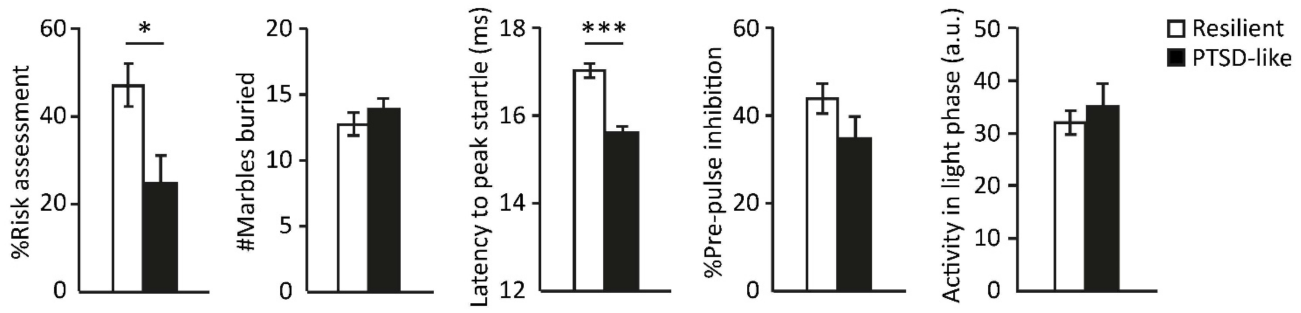


**Figure S8.** Behavioral assessment of the trauma memory cohort (experiment 3). PTSD-like animals displayed significantly reduced risk assessment behavior (t(14)=2.802, p=0.014), and shorter latencies to peak startle (t(14)=7.047, p<0.001), whereas no significant differences were observed in marble burying, percentage pre-pulse inhibition and activity during the light phase (all t(15)'s<1) from resilient mice. ***: p<0.001, *: p<0.05

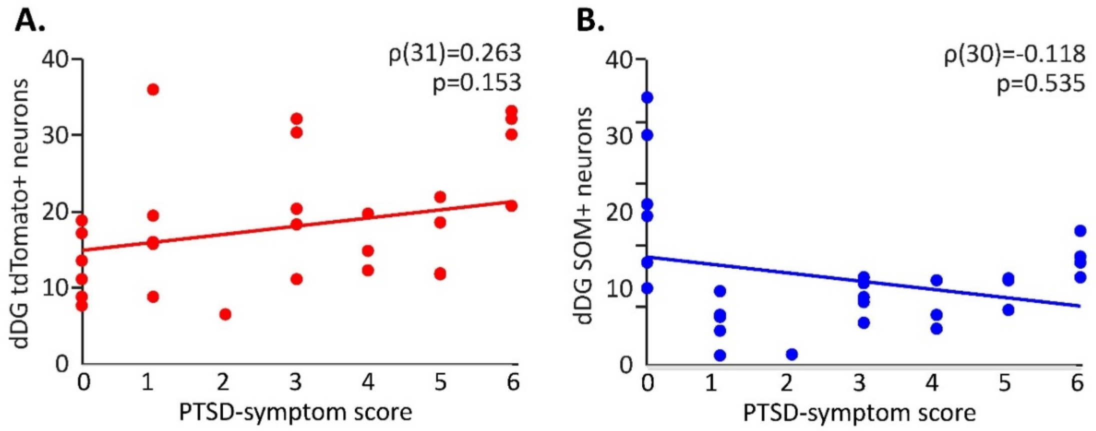


**Figure S9.** In cohort 3, neither activity during trauma memory encoding (marked by tdTomato expression) (**A**), nor the number of somatostatin (SOM) neurons (**B**) in the dorsal DG (dDG) correlated significantly with PTSD symptom score.

| | Resilient | PTSD-like |
|---|---|---|
| **Dorsal DG** | | |
| # SOM neurons active during trauma encoding | 0.00 ± 0.05 | 0.00 ± 0.71 |
| # SOM neurons active during remote trauma recall | 1.18 ± 1.60 | 0.78 ± 2.09 |
| **Ventral DG** | | |
| # SOM neurons active during trauma encoding | 0.00 ± 0.18 | 0.00 ± 0.31 |
| # SOM neurons active during remote trauma recall | 0.09 ± 2.18 | 0.45 ± 0.96 |

**Table S1.** Activity of dentate gyrus (DG) somatostatin-expressing (SOM) neurons during trauma encoding (tdTomato positive) and remote recall (c-Fos positive). Numbers represent median ± interquartile ranges.
